# Genomic profiling of *Nitrospira* species reveals ecological success of comammox *Nitrospira*

**DOI:** 10.1101/612226

**Authors:** Alejandro Palomo, Arnaud Dechesne, Barth F. Smets

## Abstract

Nitrification was long thought to consist in the stepwise oxidation of ammonia to nitrite and of nitrite to nitrate by ammonia oxidizing and nitrite oxidizing microorganisms, respectively. Recently, single microorganisms capable of complete ammonia to nitrate oxidation (comammox) were identified in the *Nitrospira* genus. This genus, previously considered to only contain canonical nitrite oxidizers is diverse and has a broad environmental distribution. Yet, a global insight into the abundance, niche preference, and genomic diversity of *Nitrospira* is missing. Here, we established the largest *Nitrospira* genome database to date, revealing 68 putative species, most without cultivated representatives. We performed a global survey through read recruitment of metagenomic data from various environments against this database that identified that environmental filtering structures species distribution, without large scale biogeographical signal. The ecological success of comammox *Nitrospira* is evident as they outnumber and are more diverse than canonical *Nitrospira* in communities from all environments but wastewater treatment plants. We detect a phylogenetic signal in *Nitrospira* species habitat preference, that is strongest for canonical *Nitrospira* species. Comammox *Nitrospira* eco-evolutionary history is more complex with subclades achieving rapid niche divergence via horizontal transfer of genes, including that encoding the hydroxylamine oxidoreductase, one of the key enzymes involved in nitrification.

## Introduction

The biological oxidation of ammonia to nitrate via nitrite, termed nitrification, is an essential process in terrestrial and aquatic environments. Nitrification links oxidized and reduced pools of inorganic nitrogen, contributes to nitrogen loss from agricultural soils reducing fertilization efficiency^1^, is a crucial process in water quality engineering^2^, and can lead to the production of nitrous oxide, a strong greenhouse gas^3^. For more than one century, this central process in the nitrogen cycle was assumed to be a two-step processes catalysed by two distinct functional groups: ammonia-oxidizing microorganism (AOM – consisting of bacteria and archaea) and the nitrite-oxidizing bacteria (NOB). In 2015, microorganisms capable of the complete oxidation of ammonia to nitrate (complete ammonia oxidation *aka* comammox) were discovered^4,5^ within the *Nitrospira* genus. This genus was previously considered to only harbour nitrite oxidizers (here also referred to as canonical Nitrospira).

Phylogenetic evidence points at a complex evolutionary history of comammox capability and it is currently unclear whether or not ancestral *Nitrospira* had this capability^6^. Clearly, the gain (or the loss) of such a trait must have had strong ecological consequences. Indeed, comammox *Nitrospira* can capture a much greater amount of energy from ammonia oxidation than canonical *Nitrospira* from nitrite oxidation, but most likely at the cost of a lower maximum growth rate^7,8^. In addition, comammox and canonical *Nitrospira* directly compete with different guilds of nitrifiers (AOM and NOB, respectively), resulting in a very different selective landscape. Yet, we have little understanding on how these markedly different metabolic strategies affect the current ecological distribution of *Nitrospira* and how such distribution relates to its evolutionary history.

Phylogenetic diversity within *Nitrospira* is high^9^: the genus consists of at least six lineages with pronounced divergence at the 16S rRNA level (sequence similarities <90%). Its members are environmentally widespread^9^, found in soils^10^, freshwater^11^, oceans^12^, groundwaters^13^, and technical systems^14,15^. Comammox *Nitrospira* has been detected in different soil and freshwater ecosystems^16^, as well as in waste and drinking water treatment systems^4,17^. Estimates from a limited number of sites indicate that comammox *Nitrospira* is abundant in drinking water treatment plants (DWTP)^18^, and to a lesser extent, in wastewater treatment plants (WWTP)^19^ and soils^20^. So far, the distribution or abundance estimates of comammox *Nitrospira* have been based on either functional gene (subunit A of the ammonia monooxygenase (*amoA*)) amplicon-based profiling^16,21^ or on limited genome-based profiles in specific environments^20,22,23^. As a consequence, our view of the distribution, abundance, and diversity of *Nitrospira* is fragmented and likely incomplete.

Shotgun metagenomic sequencing profiling has been successfully used to disclose the ecological patterns of various microbial populations at large-scale levels^24–26^. In this study, we have used metagenome assembled genomes (MAGs) to create the largest *Nitrospira* genomes database to date. We performed read recruitment analysis with metagenomes from various environments across the globe to investigate how environment and spatial distance affect *Nitrospira* distribution, and to elucidate to what degree *Nitrospira* phylogeny associates with ecology. Our analysis provides a global survey of *Nitrospira* distribution and abundance with species-level resolution and unravels the niche preferences of the different comammox *Nitrospira* types.

## Results

### *Nitrospira* database construction

Using a combination of automatic and manual binning, we retrieved 55 metagenome-assembled *Nitrospira* genomes (MAGs) from newly obtained and publicly available metagenomes (Supplementary Table S1). In addition, we downloaded 37 *Nitrospira* MAGs from public databases, resulting in a database of 92 MAGs from drinking water treatment plants (DWTP) (n = 20), freshwater (n = 30), groundwater (n = 4), soil (n = 14), wastewater treatment plants (WWTP) (n = 23) and one host-associated microbiome; from across the globe (China, n = 16; Europe, n =32; USA, = 28; Others, n = 16) (Supplementary Table S2). Average genome completeness and contamination are estimated at 85% (50% to 98%) and 2.8% (0 to 6.6%), respectively (Supplementary Table S2). The phylogenomic analysis of the 92 MAGs placed 15 MAGs into *Nitrospira* lineage I, 73 into lineage II, and 4 into other lineages (Fig. 1). These MAGs span 68 putative species (further on simply referred to as ‘species’) using a threshold average nucleotide identity (ANI) of ≥ 95%^27^ (Supplementary Fig. 1 and Supplementary Table S2). Of the 92 MAGs, 57 are comammox *Nitrospira* (32 clade A and 25 clade B) (Fig. 1 and Supplementary Table S1). Similarly, out of the 68 *Nitrospira* species 45 are comammox *Nitrospira* (25 clade A and 20 clade B) (Fig. 1).

**Fig. 1.**
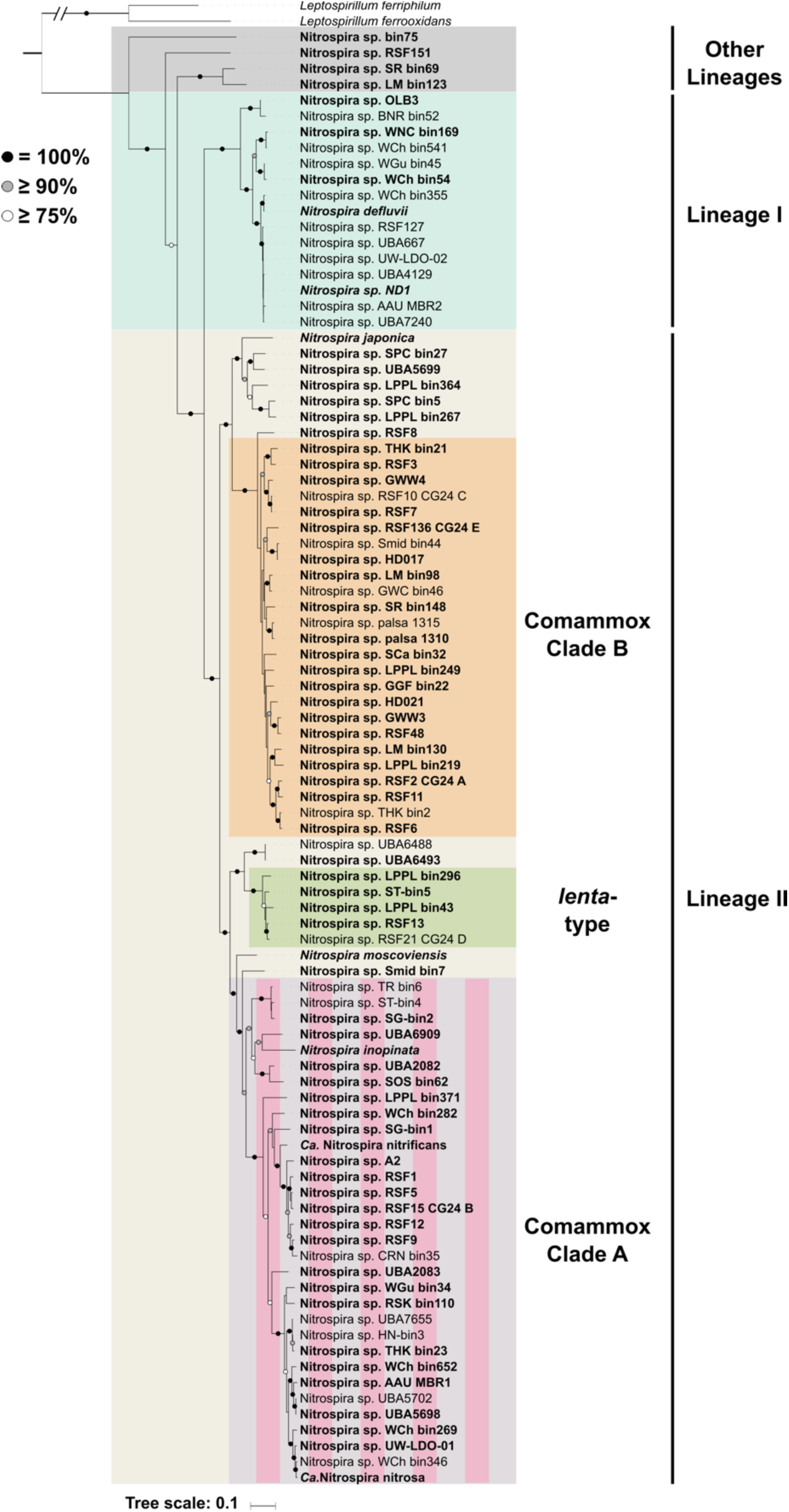
Phylogenomic affiliation of 92 *Nitrospira* MAGs inferred from the concatenation of 120 proteins. lineages and sublineages are shown with colours (lineage I, green; lineage II, yellow; Other lineages, grey; comammox clade A, pink and purple; comammox clade B, orange; lineage II lenta-type, light green). *Leptospirillum* was used to root the tree. Representative of putative *Nitrospira* species (ANI ≥ 95% are considered members of the same species) are highlighted in boldface. The strength of support for internal nodes as assessed by bootstrap replicates is indicated as coloured circles (top left legend).

### Comparison of *Nitrospira* genomes

A pangenomic analysis of 68 representative MAGs, one per *Nitrospira* species, grouped a total of 234,247 coding sequences (CDS) into 28,365 gene clusters (GCs), with a core *Nitrospira* genome consisting of 1322 GCs (Supplementary Table S3), a similar number and metabolic content to our previous study with 16 *Nitrospira* genomes^6^ (59,744 CDS grouped into 12,337 GCs, with a core genome consisting of 1382 GCs). The core genome includes genes for the nitrite oxidation pathway, the reductive tricarboxylic acid cycle for CO2 fixation (rTCA), the gluconeogenesis, the pentose phosphate, and the oxidative TCA cycle. Chlorite dismutase and copper-containing nitrite reductase (*nirK*) are also present in the core genome. 45 comammox-specific GCs were identified; 19 and 3 of these GCs have highest sequence similarity to homologs in betaproteobacterial ammonia oxidizing bacteria (AOB) and gammaproteobacterial methane oxidizers, respectively (Supplementary Table S3). These genes encode the ammonia oxidation pathway, as well as urea transporters and copper homeostasis proteins (Supplementary Table S3). We identified 61 and 31 comammox clade A and clade B-specific GCs, respectively, with mostly unknown function (Supplementary Table S3). In addition, each *Nitrospira* genome harbours an average of 272 unique gene clusters, most of unknown function.

### Comammox *Nitrospira* spp. are widely distributed

We characterized the distribution of the 68 *Nitrospira* species across 995 metagenomes across eight broadly defined environments (DWTP, n = 38; freshwater, n = 259; groundwater, n = 68; marine, n = 27; soil, n = 446; urban, n = 15; WWTP, n = 142) across the globe (China, n = 146; Europe, n =162; USA, = 502; others, n = 185) (Supplementary Table S4). *Nitrospira* species were detected in 527 metagenomes (Fig. 2). The most widely distributed *Nitrospira* species in the investigated metagenomes were the clade B comammox *Nitrospira* sp. LM_bin98 (frequency of occurrence of 25%) and the canonical lineage I *Nitrospira* sp. ND1 (21%). However, *Nitrospira* sp. LM_bin98 was in low abundance (coverage < 1) in most of the cases (88%), while *Nitrospira* sp. ND1 was detected at a higher abundance (coverage > 1) in 45% of the metagenomes where it was present (Supplementary Fig. 2). In contrast, 12 species were found in less than 1% of the metagenomes, including 4 of the 7 *Nitrospira* species with a representative isolate or enrichment (*N. inopinata, N. japonica, N. moscoviensis* and *Ca*. Nitrospira nitrificans) (Supplementary Fig. 2).

**Fig. 2.**
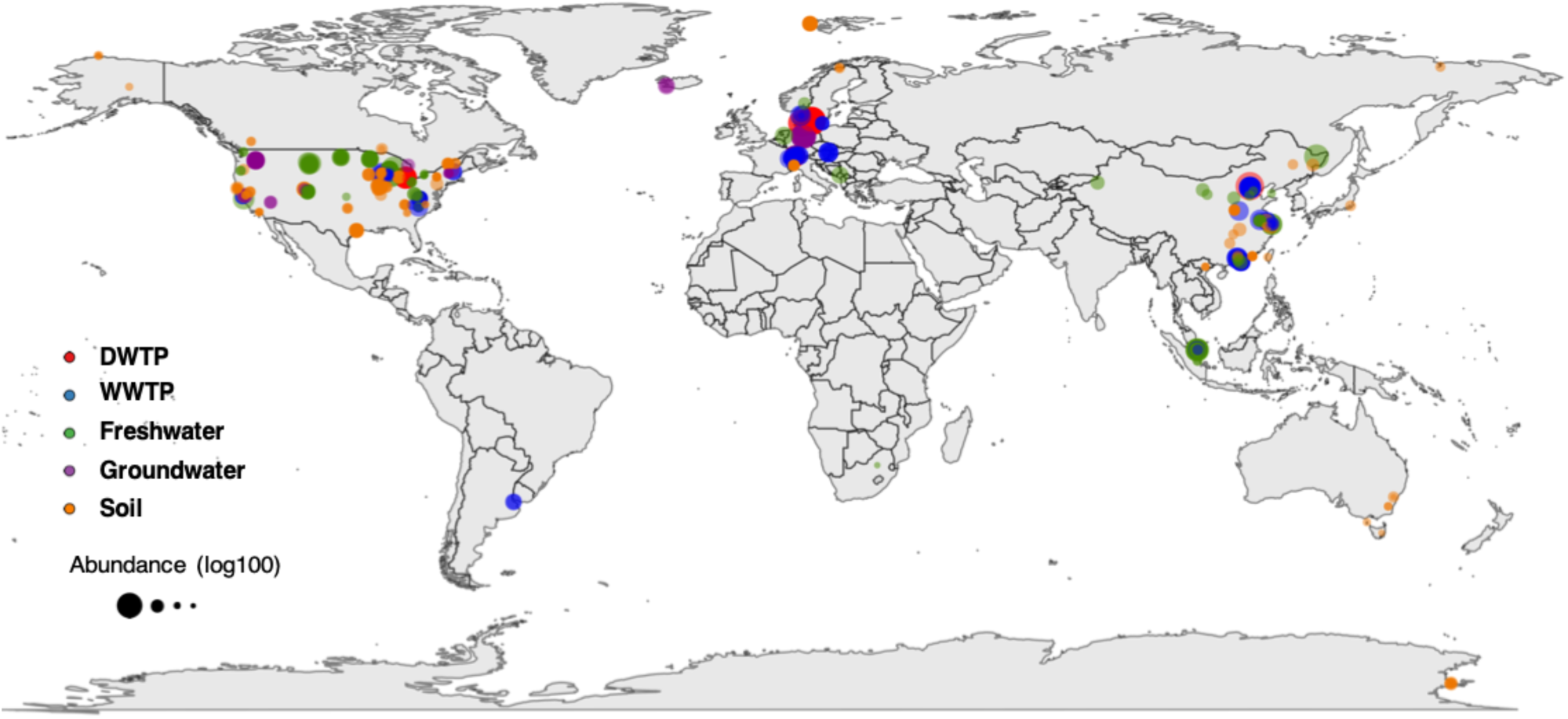
Location of the 526 metagenomes where one or more Nitrospira species were detected. The samples represent 5 distinct environments including DWTP (n = 38), WWTP (n = 130), freshwater (n = 128), groundwater (n = 39), and soil (n = 191). The size of the circle denotes the relative abundance of the Nitrospira species in the metagenome (log_100_ of the mapped reads per 20 million reads).

All the DWTP (38/38), most of the WWTP (130/147), and about half of the groundwater (39/68) and freshwater metagenomes (128/258) have at least one *Nitrospira* species. A slightly lower proportion was detected for the soil samples (191/446). On the other hand, almost none of the *Nitrospira* species in our database were encountered in marine (1/27) or built environment (0/15) metagenomes. The *Nitrospira* species richness per metagenome is highest in the DWTP metagenomes (17.5 ± 6.9, n = 38), followed by groundwater, freshwater and WWTP (groundwater = 9.4 ± 8.0, n = 39; freshwater = 6.2 ± 5.6, n = 128; WWTP= 4.8 ± 3.6, n = 130), while it is lowest in the soil metagenomes (2.0 ± 1.7, n = 191) (Fig. 3a). In all the habitats, the average number of detected comammox *Nitrospira* species exceeds that of non comammox species (*P* < 0.01), except in WWTP, where the opposite is true. The DWTP metagenomes have the highest abundance of *Nitrospira*, followed by freshwater, groundwater, and WWTP. Soil metagenomes have a very low abundance of *Nitrospira* MAGs. Comammox *Nitrospira* constitute the majority of *Nitrospira* species in all the habitats, with the exception of WWTP, where canonical species are significantly more abundant than comammox *Nitrospira* (*P* < 0.01) (Fig. 3b).

**Fig. 3.**
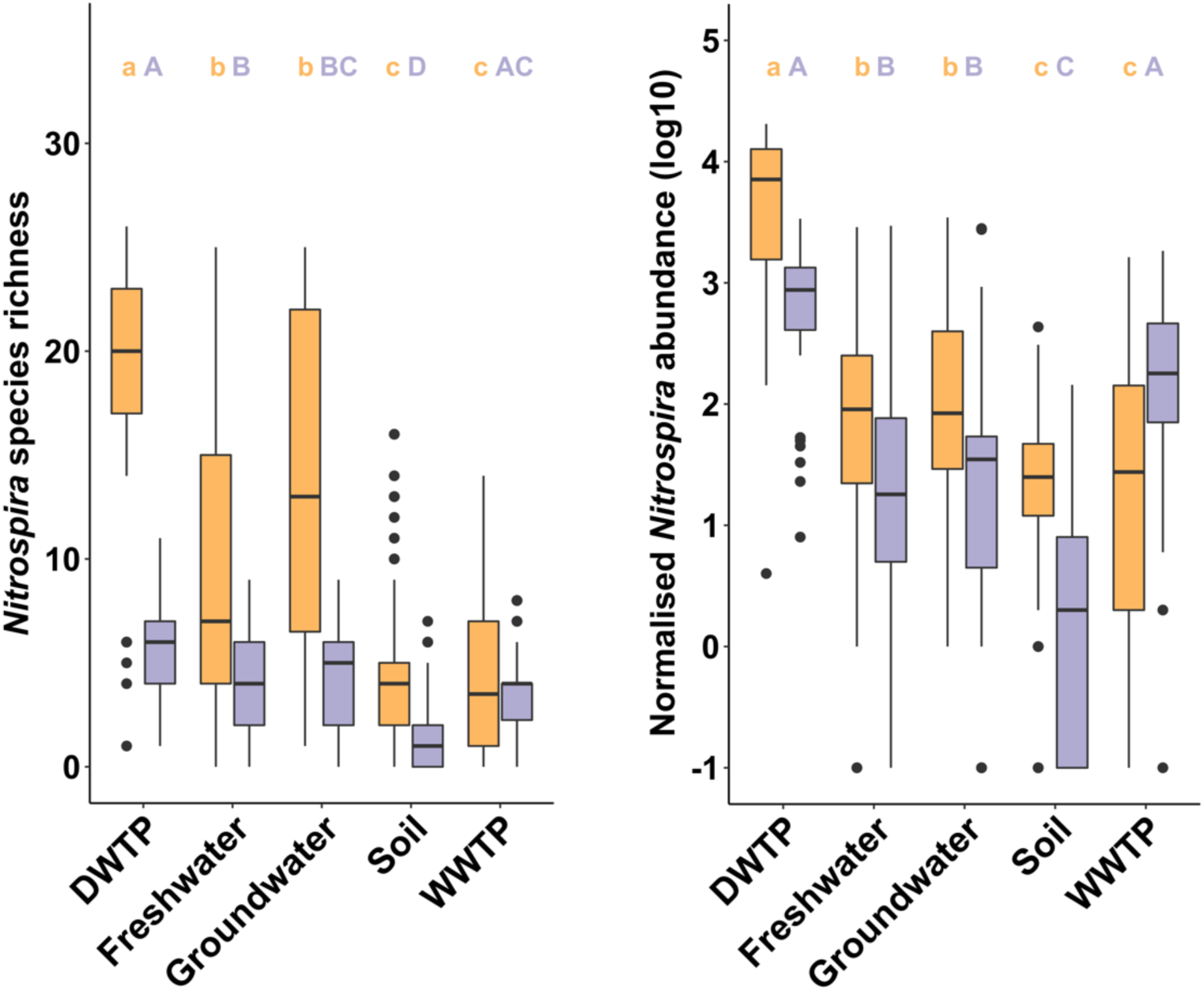
A) Richness of comammox (orange) and canonical (purple) *Nitrospira* species and B) relative abundance (mapped reads per 20 million reads, log transformed) in the different habitats. In both cases: DWTP (n = 38), WWTP (n = 130), freshwater (n = 128), groundwater (n = 39), and soil (n = 191). Differences between the mean of the richness or abundance of comammox or canonical *Nitrospira* across the habitats were assessed by a Dunn’s test; habitats with the same letter have means not significantly different from each other (p < 0.01), with small letter for comammox *Nitrospira* and capital letters for canonical *Nitrospira* species. For all habitats, the mean richness and abundance of comammox and canonical *Nitrospira* was significantly different (P < 0.01). Boxes represent the first quartile, median and third quartile of distribution values, and whiskers of 1.5 × interquartile range.

### Environment determines *Nitrospira* spp. distribution, without large scale biogeographical signal

To examine the distribution pattern of *Nitrospira* species, we conducted a principal component analysis (PCA) based on the presence and absence of *Nitrospira* species in the metagenomes. WWTP metagenomes clearly cluster and separate from DWTP metagenomes (Fig. 4a); soil metagenomes also cluster, while groundwater and freshwater metagenomes have more variable *Nitrospira* compositions (Fig. 4a). Within each habitat, we found, in most cases, a weak, but significant correlation between geographical distance and *Nitrospira* community dissimilarity (Supplementary Fig. 3). However, these correlations nearly disappear when samples within small distances were excluded from the analysis, with the exception of WWTP and DWTP (Supplementary Fig. 3). The PCA reveals niche separation between *Nitrospira* (sub)lineages. Lineage I species are primarily found in WWTP metagenomes while the distribution of lineage II species is more varied. Clade A comammox *Nitrospira* species are distributed in two clusters (referred to as clade A1 and clade A2, respectively) (Fig. 4b). Clade A1 comammox *Nitrospira* co-occur with the *Nitrospira* lineage I species in WWTP samples, while clade A2 comammox *Nitrospira* are more typical of DWTP, and some of the groundwater and freshwater metagenomes, and co-occur with most of the clade B comammox *Nitrospira* and the lineage II *Nitrospira lenta*-type species. The other clade B comammox *Nitrospira* species are more typical of the soil metagenomes (Fig. 4b). Similar association between habitat preference and species (sub)lineage was noted when examining species abundances across metagenomes (Supplementary Fig. 4). Significantly higher positive correlations were found for pairs of species from the same (sub)lineages compared to pairs from different (sub)lineages (Average Pearson’s *r*=0.48 ± 0.30, n = 248 vs 0.08 ± 0.29, n = 1026; *P* < 0.01) (Supplementary Fig. 5). The correlation was weaker between comammox clade A species (*r*= 0.19 ± 0.34, n = 153) due to the difference in the distribution of clade A1 and clade A2 species. The species abundance clade A2 positively correlates with abundance of clade B (*r*=0.36 ± 0.32, n = 144) and *N. lenta*-type genomes (*r*=0.50 ± 0.22, n = 32), while species abundance of clade A1 correlates with abundance of canonical *Nitrospira* lineage I (*r*= 0.35 ± 0.28, n = 40) (Supplementary Fig. 5). Overall, these results suggest that closely related *Nitrospira* species inhabit similar habitats with the clear exception of comammox *Nitrospira* clade A species.

**Fig. 4.**
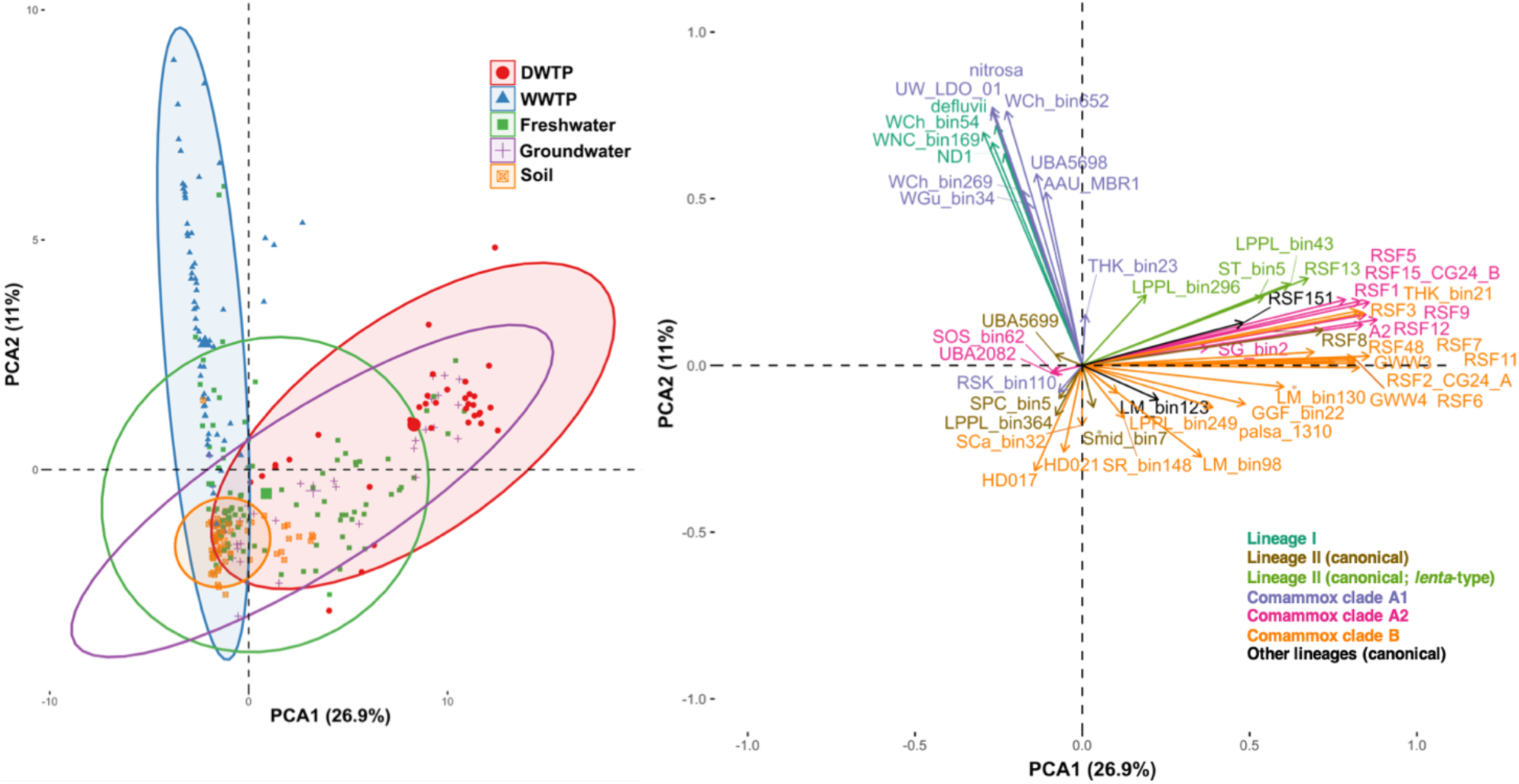
Principal component analysis based on presence or absence of *Nitrospira* species across 526 metagenomes. Left: plot of the metagenomes; 95% confidence ellipse for each habitat is also shown. Right: Plot of the *Nitrospira* species. The contributions of PC1 (horizontal axis) and PC2 (vertical axis) are 26.9% and 11%, respectively.

**Fig. 5.**
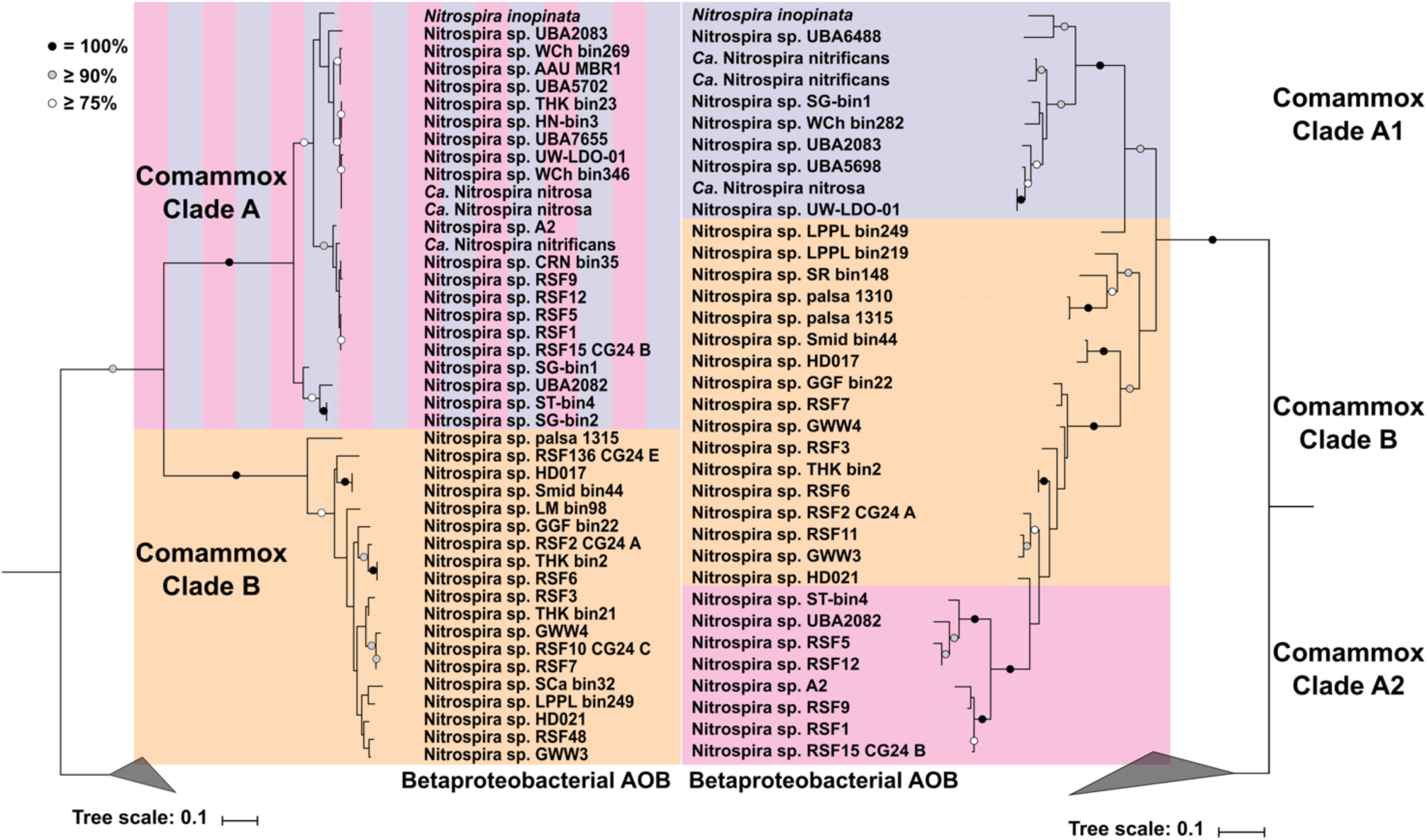
Maximum likelihood phylogenetic trees of *Nitrospira* spp. based on (left) AmoA and (right) HaoA protein sequences. Only complete or nearly complete sequences from reference species genomes are included. The strength of support for internal nodes was assessed by performing bootstrap replicates, with the obtained values shown as coloured circles (top left legend). The comammox *Nitrospira* clades are indicated by coloured boxes, while betaproteobacterial AOB sequences were used to root the trees.

### Relation between ecological and phylogenetic similarities

We assessed which feature better explained habitat preference by comparing, for pairs of species, similarity in their environmental distribution (as measured by correlation of their abundance across samples) with overall genomic similarity (as measured by ANI between their genomes), as well as, sequence similarity of key nitrification enzymes ammonia monooxygenase (AMO), hydroxylamine dehydrogenase (HAO) and nitrite oxidoreductase (NXR). Genomic similarity between species pairs positively correlates with similarity in habitat preference (Mantel statistic *r* = 0.51, P < 0.001) (Table 1). Similar results were obtained when only comammox species were evaluated (*r* = 0.56, P < 0.001) but the correlation was stronger when only the canonical *Nitrospira* species were considered (*r* = 0.78, P < 0.001). Although pairs of comammox clade A species are, on average, poorly similar in habitat preference, their genomic similarity strongly correlates with ecological similarity (Mantel statistic *r* = 0.86, P < 0.001). Thus, the heterogeneous ecological distribution of this clade can be explained by genomic dissimilarity. The opposite was observed for clade B species. Clade B species have a more homogeneous habitat preference (average Pearson *r*= 0.45 ± 0.27), but the remaining heterogeneity cannot be explained by their genomic dissimilarity (Mantel statistic *r* = 0.21, P > 0.05). Because the resolution of ANI decreases for values below 80-75%^28^ (also observed in this study; Supplementary Fig. 6), a threshold crossed for many comparison of *Nitrospira* species (Supplementary Fig. 1), we redid the analysis using a set of 120 universal single-copy proteins as metric of phylogenetic similarity (Supplementary Fig. 6). Similar trends as for ANI were observed, although with slightly weaker correlation values (Table 1). The correlation between NxrB similarity, considered a powerful functional and phylogenetic marker^9^, and habitat similarity is weaker than for whole genome similarity (especially for comammox *Nitrospira* species) (Table 1), except when only canonical *Nitrospira* species were analysed (*r* = 0.94, P < 0.01). The correlation between AmoA sequence similarity and habitat similarity is also weaker than for whole genome similarity for comammox *Nitrospira* species (*r* = 0.31, P < 0.001). HaoA similarity, however, displays a much stronger correlation with habitat similarity for all comammox species (*r* = 0.70, P < 0.001), as well as within comammox clades A (*r* = 0.76, P < 0.001) and B (*r* = 0.52, P < 0.05).

**Table 1.** Relationship (Mantel statistic r) between habitat similarity and genetic similarity between *Nitrospira* species pairs for different markers.

|  | Whole genome | Universal genes | NxrB | AmoA | HaoA |
| --- | --- | --- | --- | --- | --- |
| All *Nitrospira* | 0.51 *** | 0.39 *** | 0.31 *** | - | - |
| Canonical *Nitrospira* | 0.78 *** | 0.61 *** | 0.94 ** | - | - |
| Comammox *Nitrospira* | 0.56 *** | 0.44 *** | 0.19 * | 0.31 *** | 0.70 *** |
| Comammox *Nitrospira* - Clade A | 0.86 *** | 0.76 *** | 0.21 ** | 0.67 *** | 0.76 *** |
| Comammox *Nitrospira* - Clade B | 0.21 | 0.28 * | -0.01 | 0.33 * | 0.52 * |

\* $P < 0.05$
\*\* $P < 0.01$
\*\*\* $P < 0.001$

### Hydroxylamine dehydrogenase is potentially involved in recent niche separation within clade A

These results indicate that the hydroxylamine dehydrogenase might be a suitable indicator of niche preference for comammox *Nitrospira*. Indeed, phylogenetic analysis of HaoA shows a division of clade A into two subclusters that match those identified from habitat preference patterns (clade A1 and clade A2). Clade A2 sequences are more closely related to those of clade B species (Fig. 5). This contrasts with the clear monophyletic separation of the two comammox clades (A and B) based on phylogeny of AmoA (Fig. 6) and supported by the phylogenetic analyses of the concatenation of 120 single-copy marker proteins (Fig. 1) and of comammox *Nitrospira* specific proteins (Supplementary Fig. 7). Further, we detected two *nirK* genes (encoding a copper-containing nitrite reductase) next to the HAO cluster in all clade A2 genomes. This gene synteny is also found in several of the clade B genomes, but never in the clade A1 genomes (Supplementary Fig. 8). This shared synteny of the HAO genetic region suggests a horizontal gene transfer event between clade A2 and clade B (as also suggested by our earlier evolutionary analysis^6^) instead of convergent evolution of the hydroxylamine dehydrogenase.

## Discussion

This study represents the first effort to reveal the global distribution and ecological niches of *Nitrospira* species including comammox *Nitrospira*. We exploit large amounts of shotgun metagenomic sequencing data to quantify the abundance and relative distribution of *Nitrospira* species in different habitats through read recruitment analysis against a *Nitrospira* database of 15 universal single-copy genes^25^. The uncovered diversity of *Nitrospira* species, and specifically the vast diversity within lineage 2 *Nitrospira*, reflects the species-level diversity previously estimated from 16S rRNA sequences^9^, although it clearly underrepresents *Nitrospira* lineage IV, typical of marine environments^9^. The lack of recovered *Nitrospira* MAGs from marine metagenomes is consistent with *Nitrospira* being a minor NOB in this habitat^9^. Our analysis confirms the previously described ubiquity of *Nitrospira*^9^ and reveals the extent of the ecological success of comammox *Nitrospira*. We detected comammox species within a wide range of climatic zones, from polar (soils in Svalbard and Antarctica) to tropical (soils in Vietnam, and fresh and wastewater in Singapore) and temperate zones. We found that comammox *Nitrospira* coexist with and are more abundant than canonical *Nitrospira* in all studied habitats except in WWTP. This hitherto unrecognized dominance of comammox *Nitrospira* implies that, until recently, by equating detection of *Nitrospira* 16S rRNA or *nxrB* with strict NOB metabolism, the number of ammonia oxidizers in the environment has been systematically and significantly underestimated. Of the studied habitats, comammox *Nitrospira* is especially diverse and abundant in drinking water treatment systems, which are characterized by surface-attached microbial communities and low ammonium fluxes^15,29^, a suitable environment for high growth yield microorganisms such as comammox *Nitrospira*^7,30^. On the other hand, canonical *Nitrospira* clearly outnumber comammox *Nitrospira* in WWTP. This predominance of canonical nitrite oxidizers in WWTP is consistent with the proposition that nitrogen-rich environments select for division of labour because it maximizes growth rates^7,31^. However, in some full-scale WWTP^19,32^ and in a lab-scale low nitrification reactor^33^ comammox *Nitrospira* have been found more abundant than canonical ammonia oxidizers, suggesting that factors beyond nitrogen fluxes influence comammox *Nitrospira* fitness in these systems.

In contrast to what has been observed for other bacterial taxa such as *Streptomyces*^34^, SAR11^35^ or Acidobacteria^36^, we did not detect a strong biogeographical signal for *Nitrospira* spp., with the exception of drinking water treatment plants and, to a lesser extent, wastewater treatment plants, where dissimilarity of the *Nitrospira* communities correlates with geographical distance. However, due to the limited number of DWTP samples, a larger dataset is needed to confirm this observation. For all other habitats, the correlation nearly disappears after few tens of kilometres, which might correspond to the limit of effective dispersal of *Nitrospira* or might reflect the decline of similarity in their environmental distribution as geographical distance increases. Environmental filtering controls *Nitrospira* species distribution and, since closely related species tend to share habitats preference, it results in a clear phylogenetic clustering of *Nitrospira* types in environmental communities. As observed for other microbial taxa^37,38^, dissimilarity in habitat preference generally increases with genomic dissimilarity, with the strongest correlation for canonical *Nitrospira* species, consistent with progressive phylogenetic and ecological divergences, as expected from genetic drift or from random fluctuation in natural selection^39^. The similarity of AmoA and NxrB sequences, considered as functional and phylogenetic markers for ammonia and nitrite oxidation, respectively, do not predict niche preference for comammox *Nitrospira*. Instead, HaoA sequence similarity strongly correlates with comammox *Nitrospira* habitat preference. Following the aforementioned phylogenetic clustering, lineage I *Nitrospira* species co-occur and are mainly restricted to WWTP, while lineage 2 species are widely distributed across different habitats. Comammox clade B species are found in DWTP, groundwater, freshwater and soils. We observed a striking niche separation between closely related species of clade A comammox *Nitrospira*. While clade A2 species were mainly found in DWTP, groundwater and freshwater, sharing habitats with clade B comammox species, comammox clade A1 species primarily occupy the habitat typical of lineage I *Nitrospira* species (WWTP). Ecological divergence of closely related types can emerge from gene acquisition through horizontal gene transfer^40–42^ or by changes in existing genes whose divergence can be accelerated by natural selection^43–45^. The high sequence divergence for HAO subunits between clade A1 and clade A2 genomes, and the similarity between clade A2 HAO and clade B HAO is unlikely to result from convergent evolution, and horizontal gene transfer is the likely cause of the ecological divergence between clade A1 and clade A2 comammox *Nitrospira*. Indeed, clade A2 and some of the clade B genomes share a unique *hao* synteny that includes a duplicate *nirK*, which is absent in clade A1 genomes. Such transfer event between clade A2 and B had been proposed in a previous, more limited evolutionary analysis^6^. It is plausible that this transfer would provoke a niche modification in the recipient as HAO is one of the key enzymes for ammonia oxidation^3^. The apparent co-transfer of nearby *nirK* genes is intriguing, as NirK has recently been posited as essential in ammonia oxidation^46^. Nevertheless, although our analysis show that the genomes involved in this proposed horizontal gene transfer tend to share similar habitats, further ecophysiological analysis is needed to confirm the basis of the ecological divergence within clade A. This will require continuous efforts at isolating representatives of all comammox *Nitrospira* (sub)clades, beyond the few currently clade A1 isolate^30^ and enrichments^5^.

Taken together, our study expands the genomic inventory of the *Nitrospira* genus, exposes the ecological success of complete ammonia oxidizers within a wide range of habitats, and identifies the habitat preferences (sub)lineages of canonical and comammox *Nitrospira* spp.

## Methods

### Sample collection and DNA extraction

Filter material (15 ml) was collected from 2 locations or from 1 location at two different times at the top of the filters of 12 Danish waterworks using a 1% hypochlorite-wiped stainless-steel grab sampler. Filter material was immediately placed into cryotubes, immersed in liquid nitrogen and stored at −80°C for further analysis. DNA was extracted from 0.5 g of sand material using the MP FastDNA Spin Kit (MP Biomedicals LLC, Solon, USA) as described elsewhere^15^.

### Library preparation and sequencing

DNA libraries were generated using the 24 extracted DNA with the Nextera XT DNA library preparation kit (Illumina Inc.) according to the manufacturer’s instructions. The samples were sequenced in one lane, on a Hiseq 4000, 150bp Pair-end with dual indexing at BGIs facility in Copenhagen.

### Recovery and assessment of metagenomic assembled genomes

Trimmomatic v0.22^47^ was used to remove adapters and trim the reads (threshold quality=15; minimum length=40). Quality control was carried out using FastQC (Babraham Bioinformatics (http://www.bioinformatics.babraham.ac.uk/projects/fastqc/)). High-quality reads from each sample were co-assembled into scaffolds using IDBA-UD^48^ with the options --pre_correction --min_contig 1000. In addition, 24 single-sample assemblies were performed following the same procedure. Metagenomic binning was applied to the co-assembly using MetaBAT^49^, MaxBin2^50^, CONCOCT^51^, MyCC^52^, Binsanity^53^and COCACOLA^54^(Supplementary Fig. 9). The quality of the bins produced with the mentioned tools were improved using Binning refiner^55^ and DAS Tool^56^. In addition, metaWRAP^57^ binning and refinement modules were applied to the co-assembly. dRep^58^ was used to dereplicate all the generated bins with the secondary clustering threshold set at 99% genome-wide Average Nucleotide Identity (gANI). On the other hand, mmgenome^59^ was applied to the single-sample assemblies to recover *Nitrospira* genomes following the strategy described elsewhere^6^. The reassembly module of metaWRAP was used to improve the quality of the bins. All the resulting bins were aggregated and then dereplicated using dRep with the secondary clustering threshold set at 99% gANI. Scaffolds within the resulting dereplicated bins with divergent GC-content or tetranucleotide frequencies were removed using the RefineM^60^ with the option coverage correlation criteria (--cov_corr) set to 0.8. Furthermore, metaWRAP binning and refinement modules were applied to the co-assembly of the 6 samples used in Palomo et al.^22^. The generated bins together with the reported in Palomo et al.^6,22^ were dereplicated using dRep with the secondary clustering threshold set at 99% gANI. RefineM was applied on the resulting dereplicated bins as describe above. These bins, together with the bins recovered from the 12 waterworks of this study were dereplicated as described above. The assembly quality resulting dereplicated bins were improved by alignment against related complete or draft genomes using the Multi-Draft based Scaffolder (MeDuSa)^61^.

In addition, metagenomic binning from metagenomes downloaded from public databases (Supplementary Table S1) was carried out using the binning, refinement and reassembly modules of metaWRAP. Trimming and quality control of the raw reads as well as *de novo* assemblies of these metagenomes were carried out as describe above.

### Database construction

55 *Nitrospira* genomes recovered as part of this study together with 37 published *Nitrospira* genomes (downloaded from NCBI) were included in the genome dataset. Completeness and contamination rates of the all final population bins were assessed using CheckM^62^

For the metagenomic datasets, in addition to the 24 samples of this study and the 2 samples from Palomo et al.^22^, raw reads from 969 metagenomes downloaded from NCBI^63^ and MG-RAST^64^ were included in the analysis. This metagenomes were selected as follow: The NCBI Sequence Read Archives (SRA) (sequencing type “whole genome sequencing”, “HiSeq”, and environmental package: “air”, “aquifer”, “biofilm”, “biofilter”, “estuary”, “freshwater”, “groundwater”, “marine”, “metagenome”, “permafrost”, “rhizosphere”, “rice paddy”, “sand”, “sediment”, “soil”, “urban”, “wastewater”, “wetland”) were queried with *amoA* and *nxrB* sequences from several *Nitrospira* spp. to identify datasets most likely to contain sequences associated with *Nitrospira* spp. Metagenomic datasets identified as possessing *Nitrospira* spp. associated sequences were included to the metagenomic dataset. In addition, MG-RAST metagenomes not present in NCBI SAR, and with more than 2000 reads taxonomically annotated as *Nitrospira* were included in the metagenomic dataset. Trimming and quality control of the raw reads as well as *de novo* assemblies were carried out as describe above.

### Species abundance estimation

A 95% average nucleotide identity (ANI) cut-off was used to define species as proposed by Klappenbach et al.^65^. The *Nitrospira* genomes were dereplicated using dRep with the secondary clustering threshold set at 95% gANI. Among the genomes classified as belonging to the same species, the one with higher quality was chosen as representative genome for that species. The species abundance and coverage of each representative genome across the studied metagenomes was assessed using MIDAS^25^. Briefly, MIDAS uses reads mapped to 15 universal single-copy gene families (with ability to accurately recruit metagenomic reads to the correct species^25^) to estimate the abundance and coverage of bacterial species from a shotgun metagenome. We used the 68 *Nitrospira* species to build the database of universal-single-copy genes. For species abundance and coverage estimation the option -n was set to 20 million (1 million reads are considered sufficient to precisely estimate species relative abundance for a gut community^25^). Metagenomes were considered to contain *Nitrospira* spp. if at least five reads mapped against at least one *Nitrospira* species present in our database of 15 universal single-copy genes. For the co-occurrence analysis, only *Nitrospira* species with more than five reads mapped on more than 10 metagenomes were considered.

### Comparative genome analysis

Genomes of different *Nitrospira* species were included in the comparative genomic analysis. Gene calling was performed using Prodigal v.2.63^66^. Annotation was conducted in RAST^67^ and protein functional assignments of genes of interest were confirmed using blastp^68^. Pangenomic analysis was executed using the meta-pangenomic workflow of Anvi’o^24^ with default parameters with the exception --maxbit=0.3 (as descried in Palomo et al.^6^). Briefly, blastp was used to calculate similarities between all proteins in each genome. Weak matches between two protein sequences were eliminated using maxbit heuristic^69^. Finally, the Markov Cluster Algorithm^70^was used to generate gene clusters (GCs) from protein similarity search results. GCs were considered part of the core *Nitrospira* genome when present in at least 85% of the genomes. GCs were considered enriched on comammox *Nitrospira* when present in more than 60% of the comammox genomes (28 out of 45) and absent in more than 85% of the non comammox genomes (19 out of 21). GCs were classified as clade-specific (clade A, clade B, clade A1 and clade A2) if present in at least 55% of the clade-type genomes and absent in the rest of the *Nitrospira* genomes (Supplementary Table S3).

### Phylogenetic analysis and gene synteny

Phylogenetic analyses of *Nitrospira* genomes were conducted with the GTDB-Tk v.0.1.3 tool (https://github.com/Ecogenomics/GtdbTk) using the *de novo* workflow with a set of 120 single copy marker proteins and the genome taxonomy database (GTDB)^71^. Concatenated alignments were used to construct a maximum likelihood tree using RAxML v. 8.2.11^72^ with 200 rapid bootstraps (determined using the autoMRE option) and the LG likelihood model of amino acid substitution with gamma distributed rates and fraction of invariant sites (-m PROTGAMMAILGF; best model determined using ProtTest v. 3.4.2^73^). The tree was rooted using two *Leptospirillum* species as outgroup. The rooted tree was visualized using the online web tool from the Interactive Tree of Life (iTol)^74^. Predicted AmoA and HaoA amino-acid sequences were independently aligned with reference sequences using MUSCLE^75^. These alignments were used to construct maximum likelihood trees using RAxML v. 8.2.11 with 500 and 450 rapid bootstraps, respectively (determined using the autoMRE option). For AmoA, the tree was build using LG model plus gamma model of rates across sites considering the amino acid frequencies of the data set evolution (-m PROTGAMMALGF; best model determined using ProtTest v. 3.4.2), while for HaoA the tree was constructed using LG model with an estimation of invariable sites and gamma distribution (-m PROTGAMMAILG; best model determined using ProtTest v. 3.4.2). Both trees were rooted using two gammaproteobacterial AOB species as outgroup. The rooted trees were visualized using the online web tool from the Interactive Tree of Life (iTol)^74^.

The amino-acid sequences from the other comammox-specific proteins were independently aligned with reference sequences using MUSCLE^75^, and the maximum likelihood trees were constructed in MEGA X^76^ using the Jones Taylor Thornton model with gamma distribution and with 100 replicates of bootstrap analysis.

Gene arrangement of ammonia oxidation related genes was visualized using the R package genoPlotR^77^.

### Statistical Analyses

All statistical tests were performed using R v3.4.4^78^. Statistical significance of the mean richness and abundance of *Nitrospira* species in the different habitats were assessed using Kruskal–Wallis ANOVA followed by Dunn’s test with the Holm-Bonferroni correction.

Statistical significance of the mean richness and abundance between canonical and comammox *Nitrospira* species were evaluated using two-sided Mann-Whitney-Wilcoxon test.

Correlations between log transformed abundances of the *Nitrospira* species across 527 metagenomes were calculated using the corrplot R package^79^ with Pearson method. A Mantel test was used to investigate correlations between a matrix containing habitat similarity between species (as measured by the Pearson correlation between the abundances of *Nitrospira* species across samples) and a matrix of genome similarity (as measured by average nucleotide identity or sequence similarity of a set of 120 universal single-copy proteins between pair of genomes) or amino acid sequence similarities of key nitrification enzymes (AmoA, HaoA and NxrB). The significance of the Mantel statistic was obtained after 99,999 permutations.

*Nitrospira* community dissimilarities were calculated using the Jaccard index. The correlation between the *Nitrospira* community dissimilarities and geographic distances was calculated using the Mantel test (significance obtained after 99,999 permutations).

### Data availability

All raw sequence data and genome sequences retrieved from the Danish rapid sand filters have been deposited at NCBI under the project PRJNA384587. The rest of the retrieved draft genomes are available on figshare (http://dx.doi.org/10.6084/m9.figshare.7999448). The file containing the gene clusters sequences of the *Nitrospira* pangenome is available on figshare (http://dx.doi.org/10.6084/m9.figshare.7998641). The data that support the plots within this paper are available from corresponding author upon reasonable request.

## Acknowledgements

The authors acknowledge the contributors to the Sequence Read Achieve and MG-RAST for making their data publicly available. This study would not have been possible without this open sharing of data. The authors would like to acknowledge George Kwarteng Amoako (HOFOR), Stig Eskildsen (Forsyning Ballerup), Steffen Schulz and Jørgen Bendtsen (Nordvand), Torben Snehøj Weng (Fors A/S) and Jens Kjølby Larsen (Forsyningen) for their support in sample and data collection, as well as Florian B. Wagner and Vaibhav Diwan for providing samples from some of the waterworks. We are grateful to Neslihan Bicen and Marlene Danner Dalgaard at DTU MultiAssay Core (DMAC) Institute for Health technology for supporting library preparation and sequencing. This research was supported by a research grant (13391, Expa-N) from VILLUM FONDEN.

## Authors contributions

A.P and B.F.S conceived the study. A.P performed the bioinformatic analyses. A.P and A.D led interpretation of the results supported by B.F.S. A.P drafted the manuscript with input from A.D and B.F.S. All authors contributed to manuscript revision, and approved the final version of the manuscript.

