## Supplementary Information for "Genomic profiling of *Nitrospira* species reveals ecological success of comammox *Nitrospira*"

Alejandro Palomo[1], Arnaud Dechesne[1] and Barth F. Smets[1]
[1]Department of Environmental Engineering, Technical University of Denmark, Kgs Lyngby, Denmark

**Supplementary Information includes:**

Supplementary Figures 1 to 9

Supplementary Tables 1 to 4

Supplementary References

### Supplementary Figures

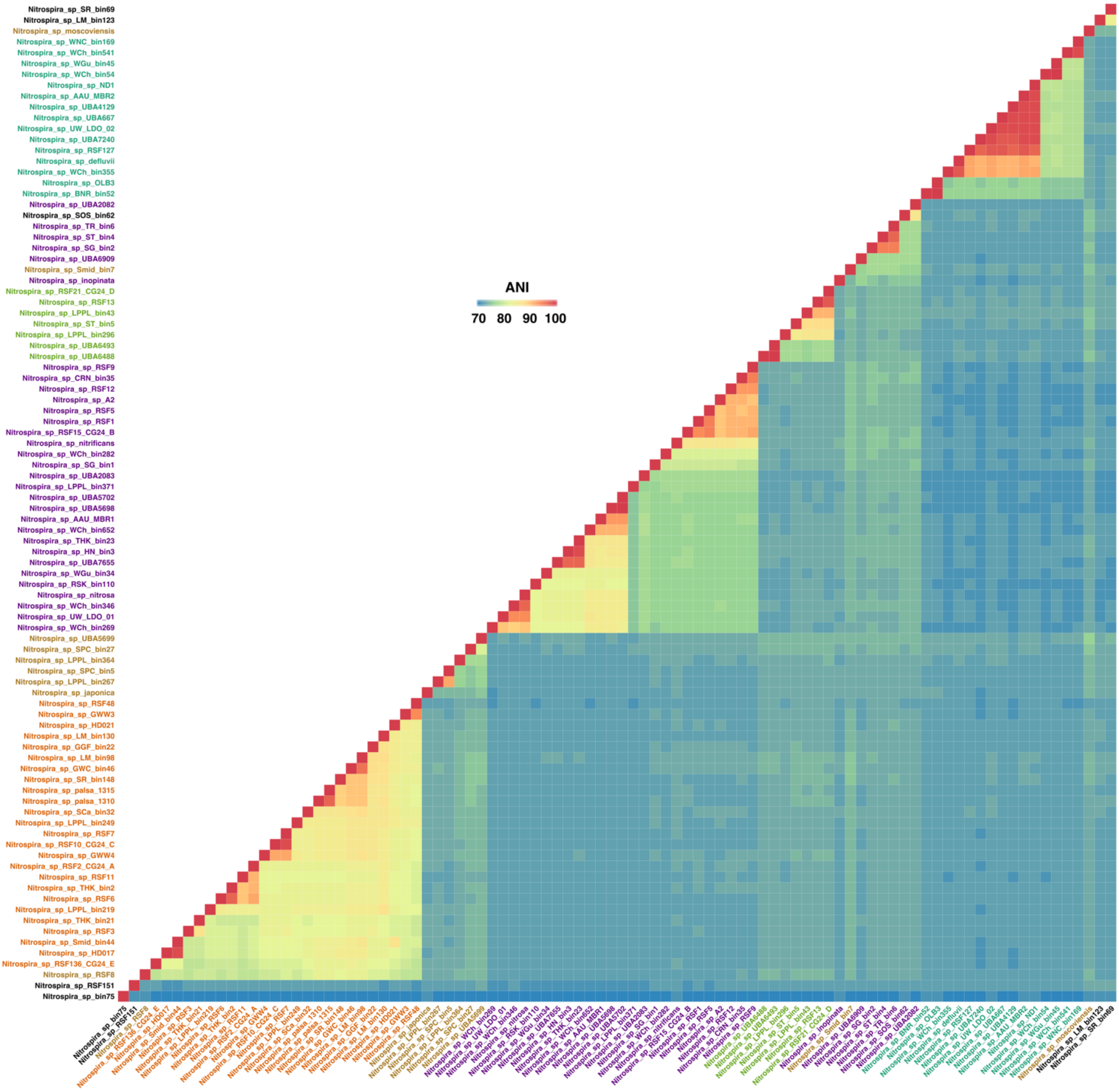


Supplementary Fig. 1. Pairwise average nucleotide identity (ANI) of pairs of *Nitrospira* genomes. The ANI was clustered using average linkage hierarchical clustering based on pairwise Euclidean distances. Colour of the *Nitrospira* genomes indicates the type (see Fig. 4).

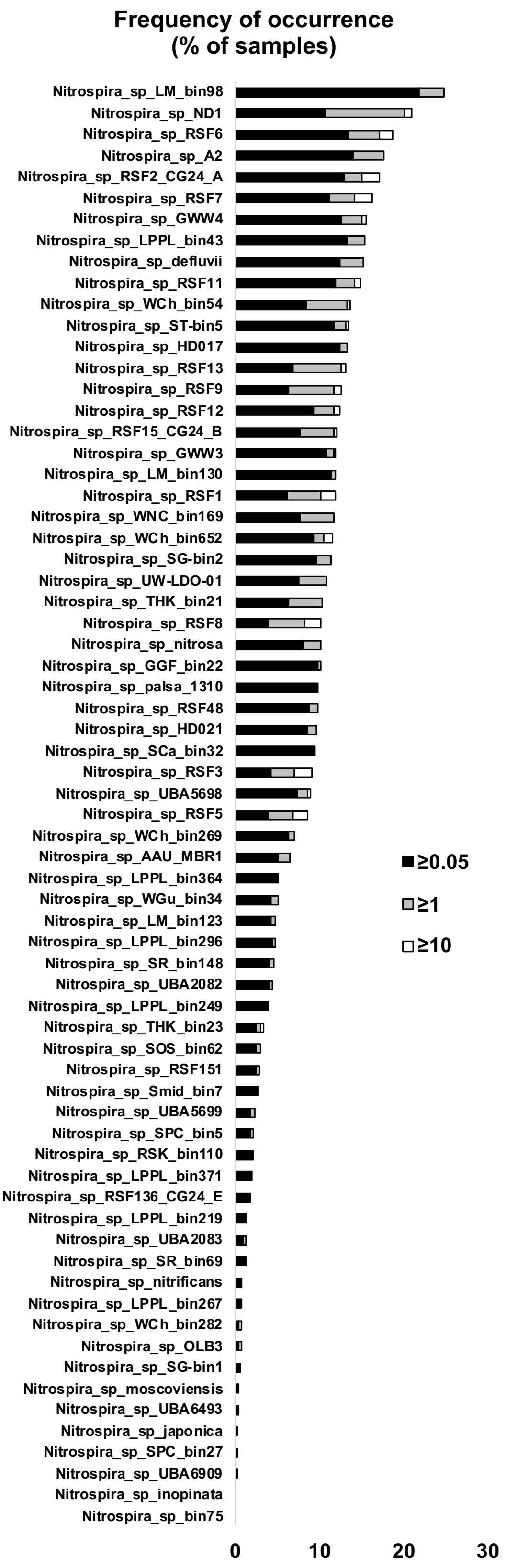


Supplementary Fig. 2. Frequency of occurrence of *Nitrospira* species in the metagenomes (n = 527) where at least one *Nitrospira* species was detected. Black, grey and white colours denote a coverage of 0.05, 1, and 10 of each species in each metagenome, respectively.

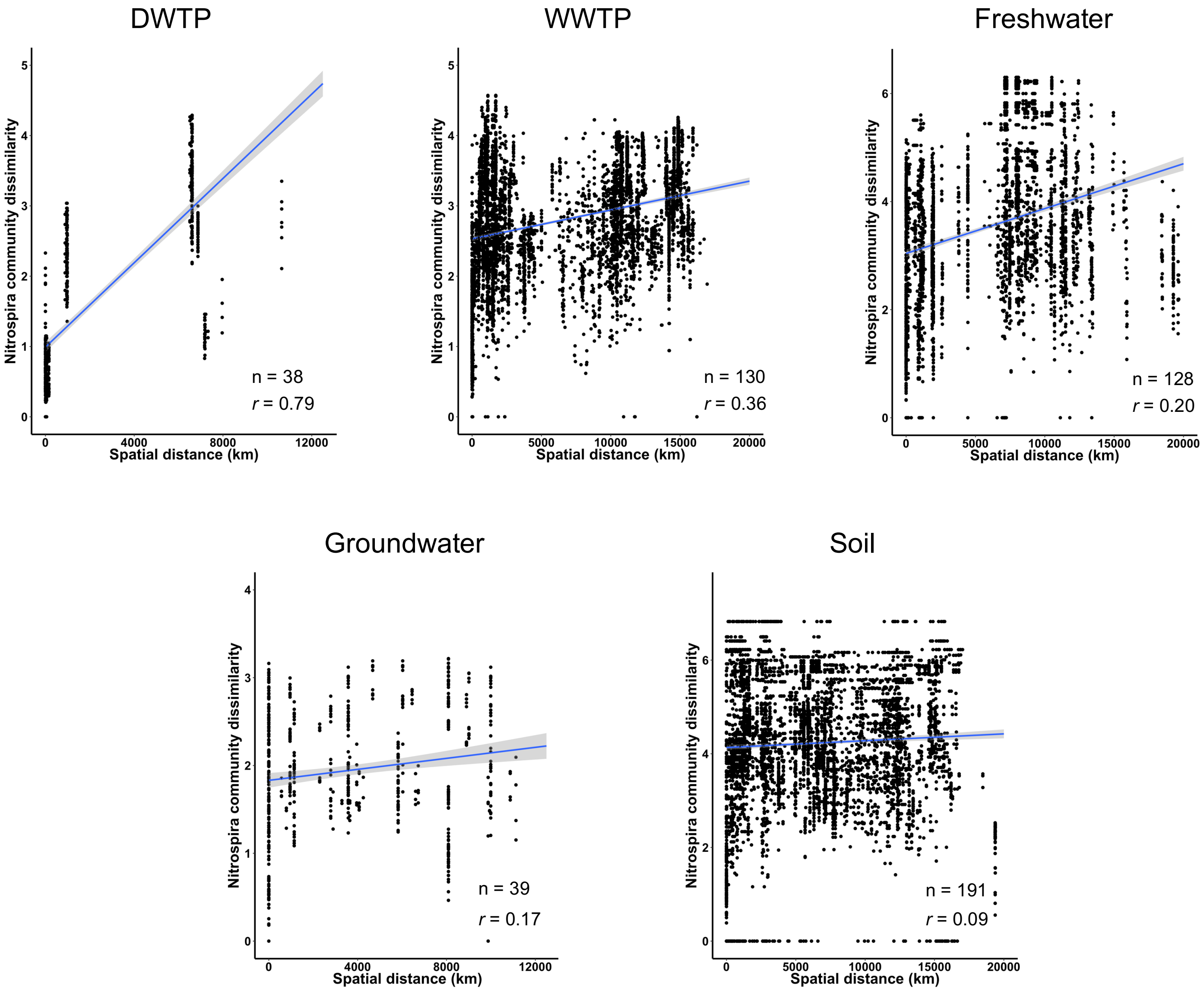


| | DWTP | Groundwater | Freshwater | Soil | WWTP |
|---|---|---|---|---|---|
| All samples | 0.79** | 0.17* | 0.20** | 0.09** | 0.36** |
| Excluding 1 km | 0.78** | 0.04 | 0.16** | 0.03 | 0.23** |
| Excluding 10 km | 0.76** | 0.04 | 0.11* | 0.01 | 0.24** |
| Excluding 50 km | 0.62** | 0.04 | 0.07* | 0.01 | 0.23** |
| Excluding 100 km | 0.58** | 0.04 | 0.06 | 0.01 | 0.21** |

* $P < 0.01$

** $P < 0.0001$

Supplementary Fig. 3. The relationship between the community similarity and the geographic distance. The dissimilarities between pairs of communities are calculated using the Jaccard index from a presence/absence matrix of *Nitrospira* genomes: the value 0 means that the two communities are the same. The Mantel test was used to test the strength and significance of correlations (*r* denotes the Mantel statistic *r*). Blue line shows the linear regression with shadowed region indicating 95% confidence intervals for the slope. The table shows the correlation (Mantel statistic r) between the community similarity and the geographic distance when all samples were analysed and when samples within short distances were excluded.

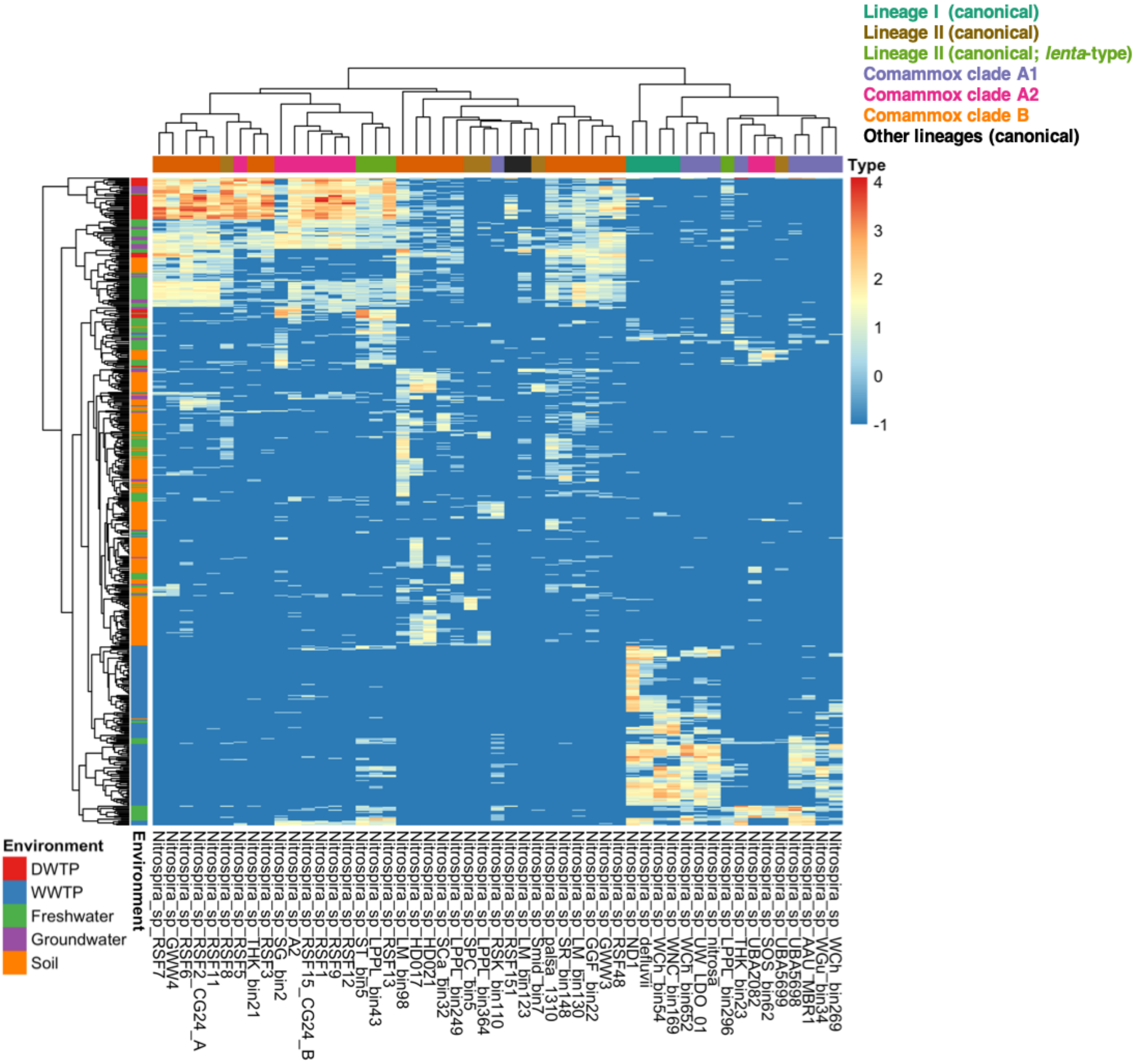


Supplementary Fig. 4. Heat map analysis of *Nitrospira* species abundances across 527 metagenomes. Dendrograms are built based on Euclidean distance. Rows represent individual metagenomes and columns represent unique *Nitrospira* species. Colour intensity indicates log transformed abundance.

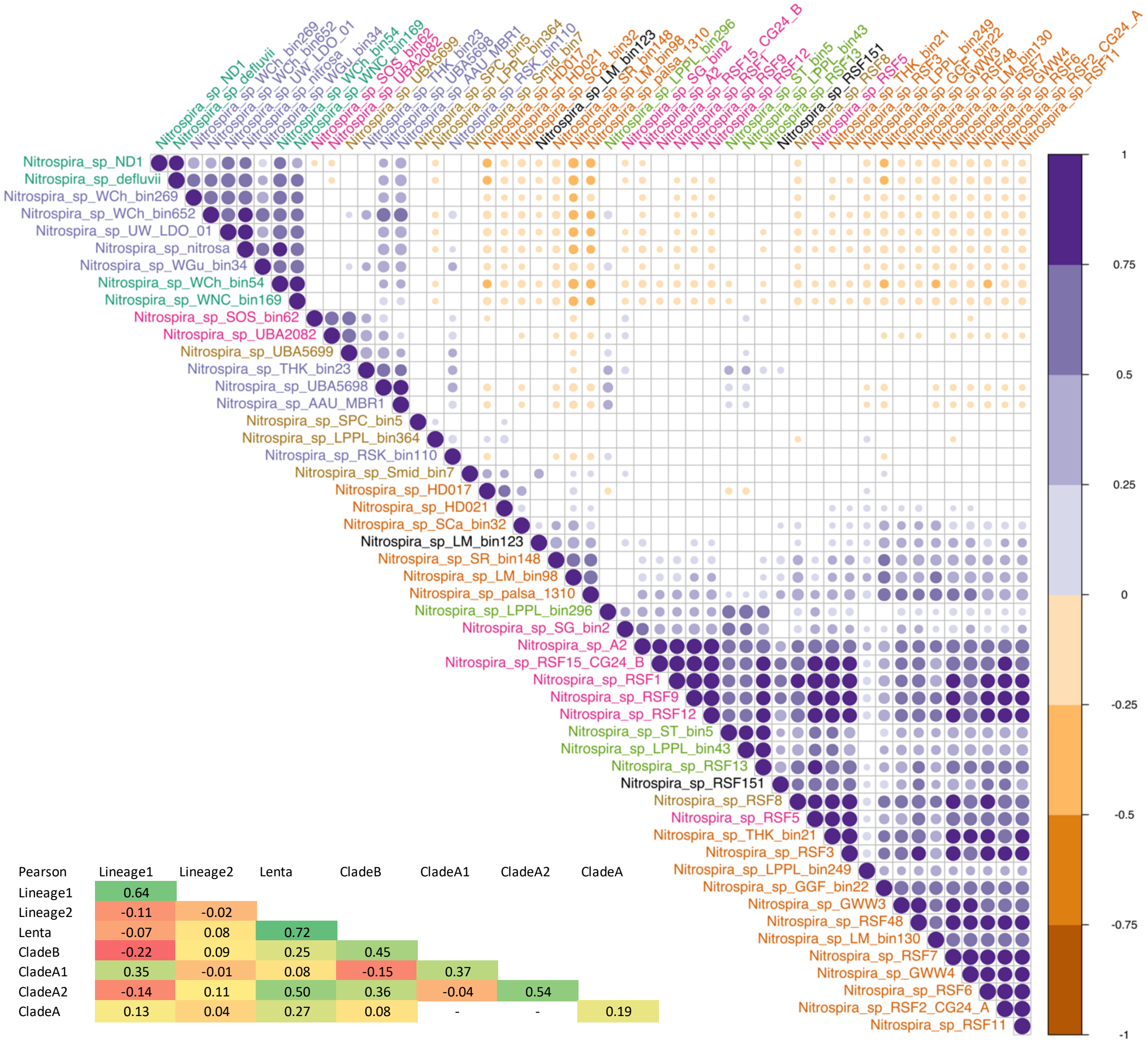


| Pearson | Lineage1 | Lineage2 | Lenta | CladeB | CladeA1 | CladeA2 | CladeA |
|---|---|---|---|---|---|---|---|
| Lineage1 | 0.64 | | | | | | |
| Lineage2 | -0.11 | -0.02 | | | | | |
| Lenta | -0.07 | 0.08 | 0.72 | | | | |
| CladeB | -0.22 | 0.09 | 0.25 | 0.45 | | | |
| CladeA1 | 0.35 | -0.01 | 0.08 | -0.15 | 0.37 | | |
| CladeA2 | -0.14 | 0.11 | 0.50 | 0.36 | -0.04 | 0.54 | |
| CladeA | 0.13 | 0.04 | 0.27 | 0.08 | - | - | 0.19 |

Supplementary Fig. 5. Correlogram showing significant Pearson correlations (α = 0.01) for log transformed abundances of the *Nitrospira* species across 527 metagenomes. Colour indicates whether the correlation is positive (purple) or negative (brown). Size and darkness of the circles indicate the strength of the correlations, with stronger correlations being larger and darker than weaker ones. Correlations with p-value > 0.01 are considered as insignificant, and omitted. Colour of the *Nitrospira* apecies indicates the type (see Fig. 4). The heatmap (bottom left) shows the average pairwise Pearson correlation of *Nitrospira* (sub)lineages.

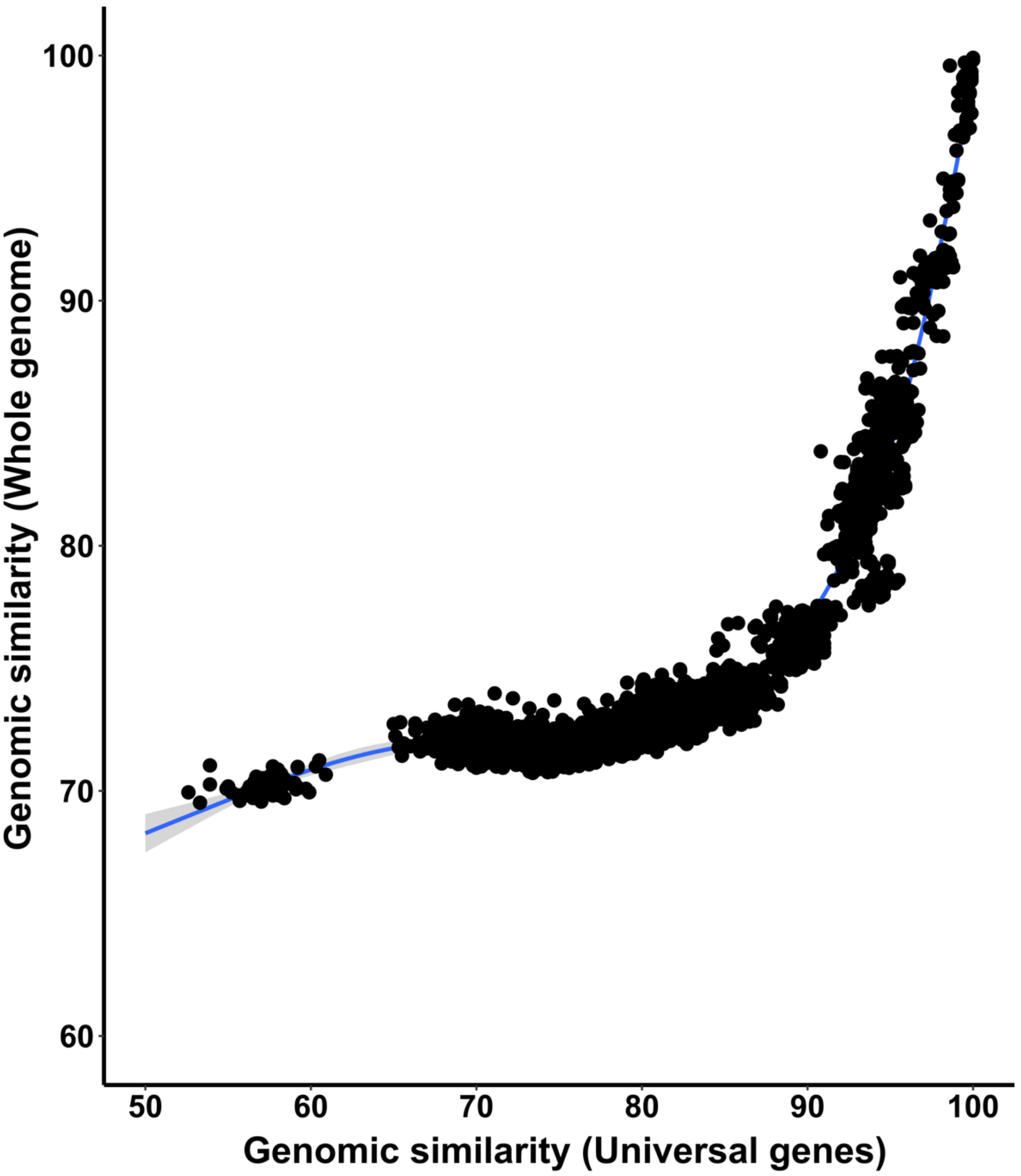


Supplementary Fig. 6. Relationship between genetic similarity based on the amino acid sequence of a set of 120 universal single-copy genes and the ANI of the whole genome of *Nitrospira* pair of species.

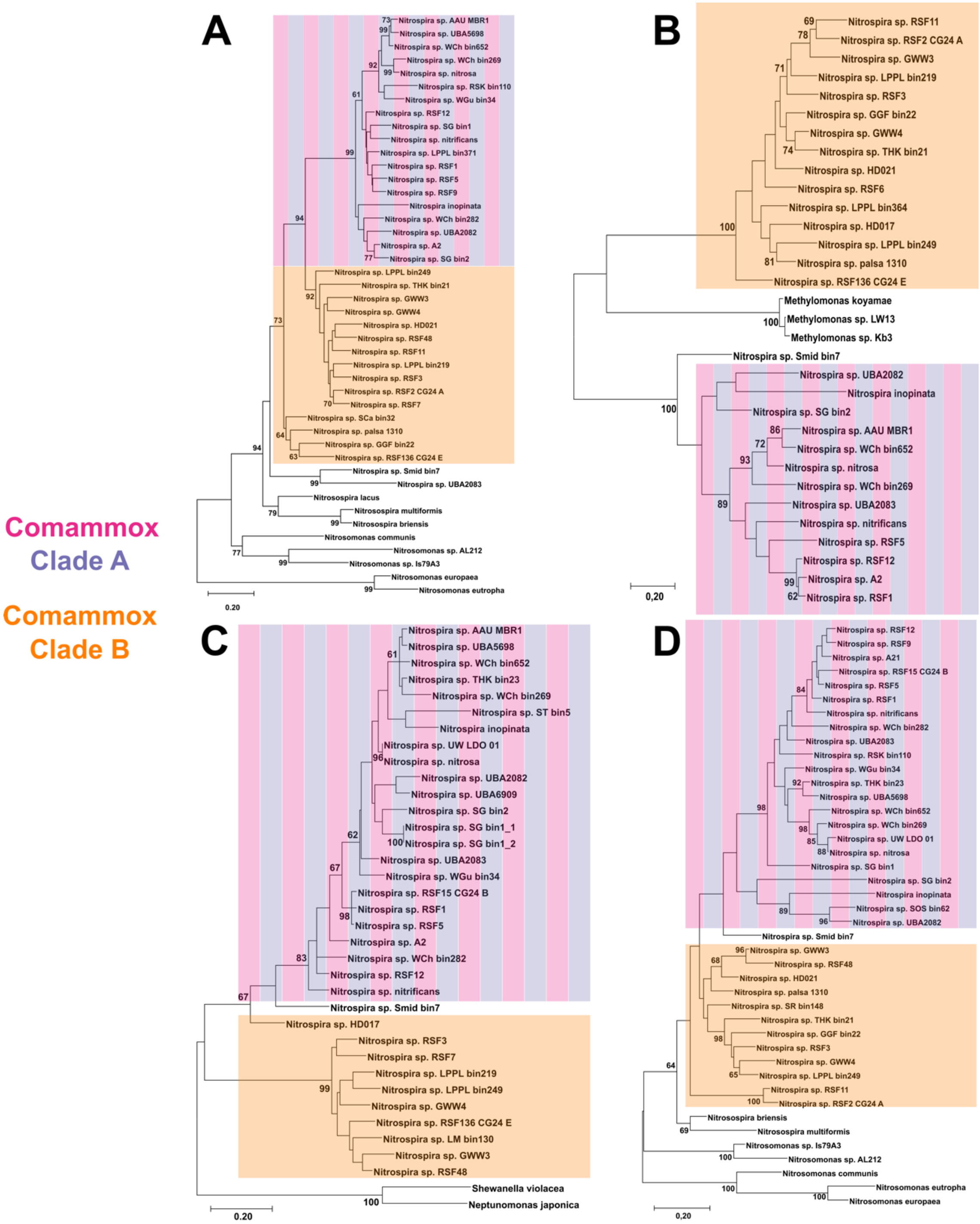


Supplementary Fig. 7. Maximum likelihood phylogenetic trees of different comammox-specific proteins: A) Urea carboxylase-related amino acid permease (GC_00001893). B) Polyhydroxybutyrate (PHB) depolymerase (GC_00002364). C) 2/2 hemoglobin type II (GC_00002249). D) Glycosyltransferase (GC_00002243). Bootstrap values greater than 60 are shown. Comammox *Nitrospira* clades are indicated by coloured boxes.

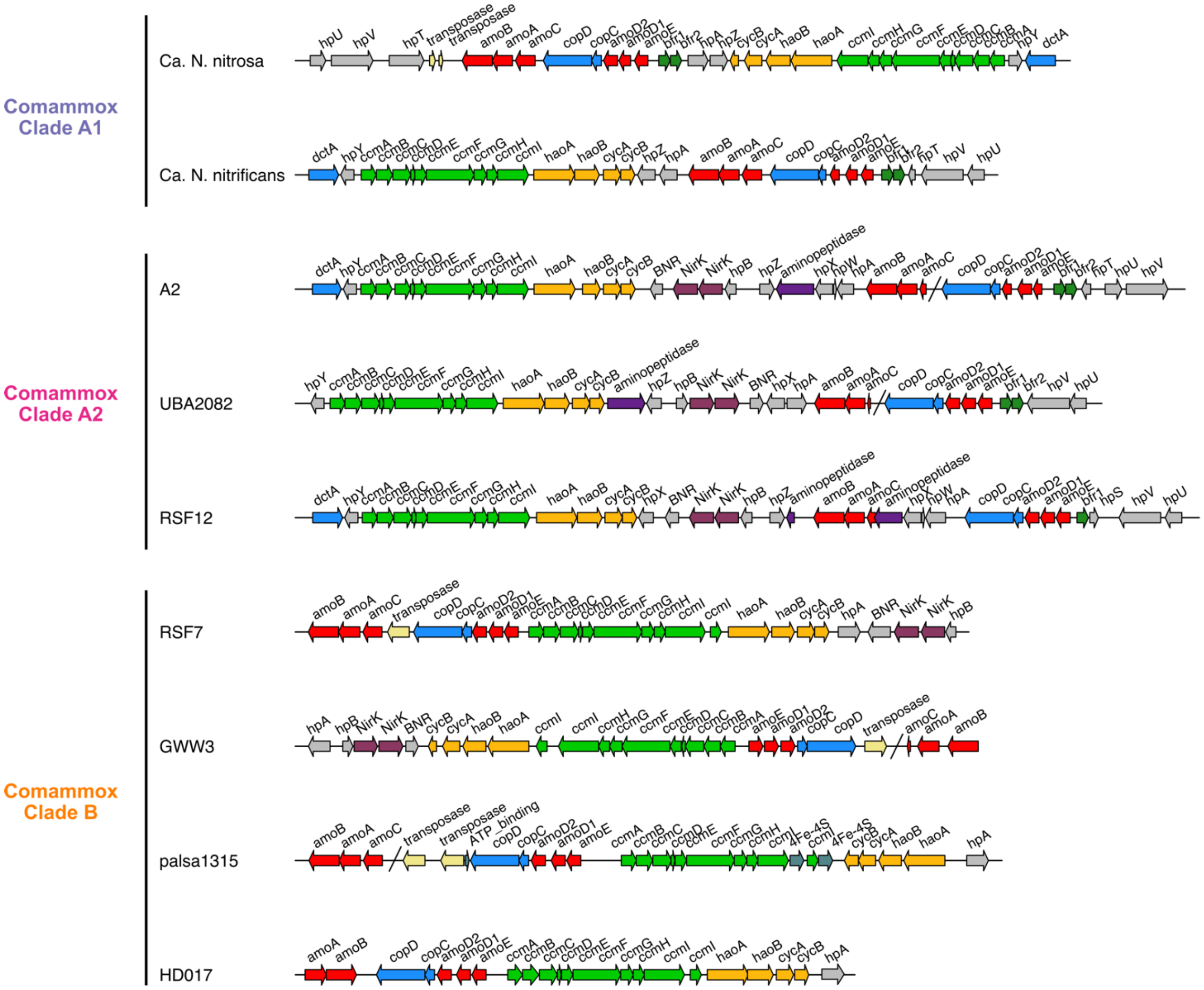


Supplementary Fig. 8. Unique shared synteny between clade A2 and some of the clade B genomes in hydroxylamine reductase genomic region (yellow arrows, duplicated NirK). Schematic of the ammonia oxidation pathway genomic region in comammox *Nitrospira* clade A1, clade A2, and clade B genomes. Functions of the encoded proteins are represented by colour. Single diagonal line designates a break due to contig fragmentation. Arrows represent genes, arrow direction denotes the orientation of the coding strand, and arrow lengths are proportional to the gene lengths. The scale bar corresponds to 5 kb sequence length.

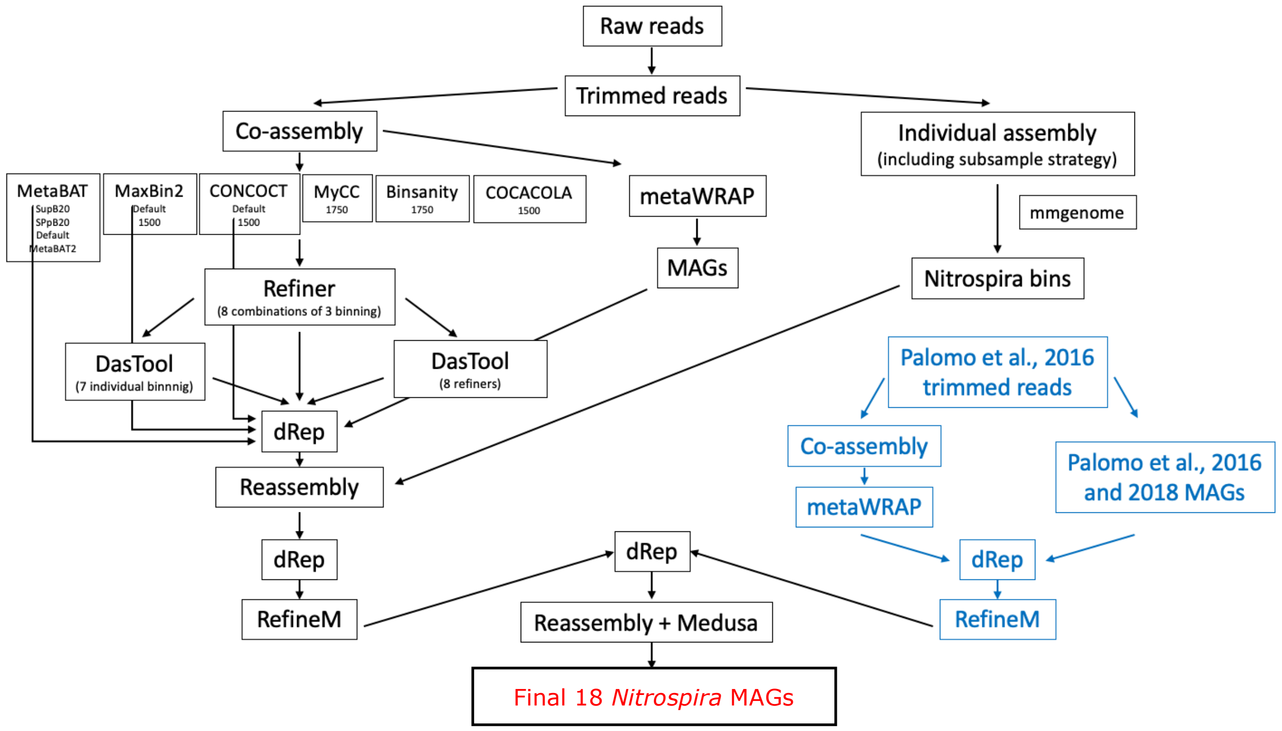


Supplementary Fig. 9. The implemented workflow for the MAGs recovery from 13 Danish groundwater-fed rapid sand filters.

#### Supplementary Tables

**Supplementary Table 1**. List of publicly available metagenomes used to retrieve metagenome-assembled *Nitrospira* genomes.
(provided as an Excel table)

**Supplementary Table 2**. Characteristics of *Nitrospira* genomes.
(provided as an Excel table)

**Supplementary Table 3**. List of gene clusters of the *Nitrospira* pangenome.
(provided as an Excel table)

**Supplementary Table 4**. Characteristics of metagenomes.
(provided as an Excel table)
